# Histone deacetylase inhibition mitigates cognitive deficits and astrocyte dysfunction induced by Aβ oligomers

**DOI:** 10.1101/2023.10.25.564038

**Authors:** Juliana Morgado, Luan Pereira Diniz, Ana Paula Bergamo Araujo, Leticia Maria da Silva Antônio, Hanna Paola Mota Araujo, Pedro de Sena Murteira Pinheiro, Fernanda Savacini Sagrillo, Gabriele Vargas Cesar, Sérgio T. Ferreira, Cláudia Pinto Figueiredo, Carlos Alberto Manssour Fraga, Flávia Carvalho Alcantara Gomes

**Author notes:** Corresponding author: Dr. Flávia Carvalho Alcantara Gomes Instituto de Ciências Biomédicas Universidade Federal do Rio de Janeiro Centro de Ciências da Saúde, Ilha do Fundão 21941-902 - Rio de Janeiro, RJ, Brazil. Those authors equally contributed to the work (Juliana Morgado, Luan Pereira Diniz, Ana Paula Bergamo Araujo).

## Abstract

Inhibitors of histone deacetylases (iHDACs) are promising drugs for neurodegenerative diseases. We have evaluated the therapeutic potential of the new iHDAC6 LASSBio-1911 in Aβ oligomer (AβO) toxicity models and astrocytes, key players in neuroinflammation and Alzheimer’s disease (AD). Astrocyte phenotype and synapse density were evaluated by flow cytometry, Western blotting, immunofluorescence and qPCR, in vitro and in mice. Cognitive function was evaluated by behavioural assays using a mouse model of intracerebroventricular infusion of AβO. LASSBio-1911 modulates reactivity and synaptogenic potential of cultured astrocytes and improves synaptic markers in cultured neurons and in mice. It prevents AβO-triggered astrocytic reactivity in mice and enhances the neuroprotective potential of astrocytes. LASSBio-1911 improves behavioural performance and rescues synaptic and memory function in AβO-infused mice. These results contribute to unveiling the mechanisms underlying astrocyte role in AD and provide the rationale for using astrocytes as targets to new drugs for AD.

## 1. INTRODUCTION

Histone acetylation/deacetylation is a key epigenetic modification that controls gene expression by modifying the chromatin structure. The histone deacetylase (HDAC) family of enzymes controls the acetylation status of several cytosolic and nuclear proteins by removing acetyl groups from the acetylated ε-amino groups of lysine residues [1]. Levels of acetylation have been reported to play a major role in the pathogenesis of several neurodegenerative diseases, such as Alzheimer’s, Parkinson’s, and Huntington’s disease. Recently, inhibitors of HDAC (iHDACs) have emerged as promising targets for neurodegenerative diseases [2–4]. Epigenetic modifications caused by the inhibition of HDAC have been associated with memory restoration and reduced cognitive deficits in several amyloidosis animal models [5, 6]. HDAC6 plays an important role in the clearance of misfolded proteins [7], control of axonal transport of mitochondria [8] and acetylation of tau [9]. While paninhibition of HDAC may affect a range of cellular functions, resulting in a nonspecific effect, selective inhibitors of HDAC6 have proven to be more specific in their actions, thus emerging as a promising candidate for therapy of neurodegenerative diseases (Simões-Pires et al. 2013; Rodrigues et al., 2020).

Alzheimer’s disease (AD) is the most prevalent form of dementia in elderly individuals, marked by progressive synaptic and cognitive dysfunction. Brain accumulation of both insoluble and soluble amyloid-β species, brain inflammation, impaired axonal transport, increased formation of neuronal reactive oxygen species, increased tau phosphorylation and formation of neurofibrillary tangles, and mitochondrial damage are all characteristic of AD pathologies [10, 11]. There is no effective treatment for AD, which may be attributed in part to a lack of a clear understanding of the underlying molecular mechanisms of disease progression. We and others have shown that defective astroglial function and reactivity significantly contribute to the development of AD [12–16] and aged-related brain dysfunction [17, 18]. Data from different mouse models of AD and single-cell seq analysis experiments in humans and mice suggest that astrocyte reactivity is highly heterogeneous and may depend, in addition to other factors, on the proximity of the β-amyloid plaques and the stage of the disease [19–21]. We previously showed atrophy of astrocyte processes and impaired synaptogenic and neuroprotective function in astrocytes after exposure to soluble amyloid-β oligomers (AβO) [14]. Rescue of these properties mitigates synaptic and memory loss induced by AβOs in mice [14], supporting the association between astrocyte reactivity and brain dysfunction induced by AβO. Recently, a neurotoxic astrocyte phenotype has also been shown to play a role in different pathological models of AD [22].

Within this context, the modulation of astrocyte reactivity emerges as an important alternative mechanism for the treatment of neurodegenerative diseases. Activation of HDACs triggers inflammatory pathways and reorganization of the astrocytic cytoskeleton, contributing to neuroinflammation [23, 24], while HDAC inhibition improves astrocyte neurotrophic and neuroprotective capacity [25]. Despite those data, there is no clear evidence on the mechanisms underlying the effects of these inhibitors on astrocytes. Therefore, the aim of this study was to evaluate the therapeutic potential of the HDAC6 inhibitor LASSBio-1911 *in vitro* and *in vivo* in an AβO toxicity model and to investigate the cellular mechanisms underlying these events, with a particular focus on astrocyte biology.

## 2. METHODS

### 2.1 Synthesis of the drugs

LASSBio-1911, (*E*)-*N*’-(4-(hydroxycarbamoyl)benzylidene)-4-(dimethylamino) benzohydrazide, was synthesized according to the methodology previously described by Rodrigues et al., 2016 [20]. After purification by column chromatography and complete characterization by using different spectrometric and spectroscopic techniques, the obtained samples were subjected to reversed-phase HPLC analysis, which indicated a purity > 95%.

### 2.2. Methodology of docking analysis

The protein data bank high-resolution crystal structure 5EEN (1.86 Å) of HDAC6 catalytic domain 2 in complex with belinostat (DOI: 10.1038/nchembio.2134) was selected for the docking studies. The analyses were performed using the GOLD v.2021.2 program (CCDC). The four fitness functions available in the program, namely, ASP, [26] ChemPLP, [27, 28] ChemScore and GoldScore, [29] were evaluated in the redocking analysis of the cocrystallized ligand to identify the most adequate fitness function. Crystallographic water molecules were removed during the docking runs, and the binding site was determined within a 10 Å distance from the native ligand. In this way, the obtained root-mean square deviation (RMSD) values between the best result for each fitness function and the crystallographic structure indicated ChemScore as the best function to proceed with. The higher performance of the ChemScore function was related to the correct identification of the chelating interaction between the ligand’s hydroxamic acid moiety and the zinc ion in a bidentate pattern. ChemScore was used for the docking study of LASSBio-1911 into HDAC6 catalytic domain 2 (5EEN).

### 2.3 Preparation and characterization of Aβ oligomers (AβO)

AβO were prepared weekly from synthetic Aβ_1-42_ (American Peptide Company, Sunnyvale, CA) and were routinely characterized by size-exclusion chromatography and, occasionally, by western immunoblots, as previously described [30–32] AβO ranged from dimers (∼9 kDa) to higher molecular weight oligomers (∼50-100 kDa) [33]. Oligomer preparations were aliquoted and kept at −70 °C until use.

### 2.4 Animals

Newborn (P0) Swiss mice were used for astrocyte cultures, and embryonic 14-day-old mice (E14) were used for neuronal cultures. For *in vivo* experiments, we used 2- to 3-month-old male Swiss mice. All animals were housed under standard conditions with ad libitum access to food and water. Animal handling and experimental procedures were previously approved by the Animal Use Ethics Committee of the Federal University of Rio de Janeiro (CEUA-UFRJ, approval protocol 006/18). Experiments were performed according to Brazilian Guidelines on Care and Use of Animals for Scientific and Teaching Purposes (DBCA). Animals used in this study were derived from the animal facility of the Instituto de Ciências Biomédicas (ICB/UFRJ) and Faculdade de Farmácia (UFRJ).

### 2.5 Neural cell cultures and treatments

Primary astrocyte cultures were derived from the hippocampus of newborn Swiss mice as previously described [34]. Briefly, the hippocampus was removed, and the meninges were carefully stripped off. Tissues were maintained in Dulbecco’s minimum essential medium (DMEM) and nutrient mixture F12 (DMEM/F12, Invitrogen) supplemented with 10% foetal bovine serum (FBS, Invitrogen). Cultures were incubated at 37 °C in a humidified 5% CO_2_, 95% air chamber for approximately 7-10 days *in vitro* (DIV) until confluence.

Neuronal cultures were prepared from embryonic Day 15-16 Swiss mice. Briefly, the hippocampus was removed, meninges were carefully removed, neural tissue was dissociated in neurobasal medium (Invitrogen), and the cells were plated at a density of 75,000 per well of 13 mm diameter onto glass coverslips previously coated with poly-L-lysine (10 µg/mL, Sigma). Cultures were maintained in neurobasal medium supplemented with B-27, penicillin, streptomycin, fungizone, L-glutamine, and cytosine arabinoside (0.65 µM, Sigma) at 37 °C in a humidified 5% CO_2_, 95% air atmosphere for 12 DIV. Purified cultures of neurons and astrocytes were treated with 1 μM LASSBio-1911 for 24 hours, after which the cells were processed for morphological or biochemical assays.

### 2.6 Astrocyte conditioned medium preparation and neuronal treatment

To obtain conditioned medium from LASSBio-1911-exposed astrocytes (CM 1911), confluent astrocyte cultures were kept for 4 hours in serum-free medium, after which they were exposed to 1 μM LASSBio-1911 or vehicle for 24 hours (for CM 1911 or CM, respectively). Cells were washed three times to eliminate drug, and fresh serum-free DMEM/F12 medium was added. The medium was collected after 24 hours and centrifuged at 1,500 × g for 10 min to remove cellular debris. To test the effect of astrocyte-conditioned medium, neuronal cultures were previously treated at 37 °C for 30 min with CM or CM 1911 and then exposed to 500 nM AβO or an equivalent volume of vehicle (2% DMSO in PBS) for 3 hours.

### 2.7 MTT

After 24 hours of treatment, the cultures had their media collected for subsequent assays, and the cells were incubated at 37 °C in a humidified oven for 2 hours with MTT (Sigma‒ Aldrich, 5 µg/ml). The medium was removed, and 100% DMSO (Sigma‒Aldrich) was added to each well to dissolve the formazan salts. The plates were read at 540 nm in a plate reader (Molecular Devices).

### 2.8 Nitrite level measurement

Nitric oxide (NO) production was determined indirectly through a nitrite (NO_2_) assay. NO_2_ is stable metabolite of NO, according to the Griess reaction [35]. Briefly, a 50 µL aliquot of media was mixed with an equal volume of Griess reagent [0.1% N-(1-naphthyl) ethylenediamine dihydrochloride, 1% sulfanilamide, and 2.5% phosphoric acid] and incubated at 22 °C for 10 min, followed by absorbance measurement at 540 nm. Based on a standard curve of NaNO_2_ (Sigma‒Aldrich) ranging from 0 to 100 µM, the nitrite concentration was calculated. Background NO_2_^-^ was subtracted from each experimental value.

### 2.9 Immunocytochemistry

Astrocyte and neuronal cultures were fixed with 4% PFA in PBS (pH 7.4) for 15 min, and nonspecific sites were blocked with 3% bovine serum albumin (BSA; Sigma‒Aldrich), 5% normal goat serum (Sigma‒Aldrich) and 0.2% Triton X-100 diluted in PBS for 1 h before incubation with the following antibodies: mouse anti-GFAP (1:1,000; Millipore, Cat. MAB360), rabbit anti-LCN-2 (1:200; Millipore, Cat. AB2267), rabbit anti-GFAP (1:1,000; DAKO Cytomation, Cat. Z0334), rabbit anti-Spinophilin (1:500; Abcam, Cat. Ab18561), mouse anti-Synaptophysin (1:1,000; Millipore, Cat. MABN1193), mouse anti-Histone H3 (1:200-1:400; Cell Signalling, Cat. #14269), rabbit anti-acetyl histone H3 (1:500; Cell Signalling, Cat. #9677), rabbit anti-C3 (1:200; Abcam, Cat. ab11887), mouse anti-S100a10 (1:500; Abcam, Cat. ab232524) at 4 °C overnight. Subsequently, the cells were thoroughly washed with PBS and incubated with secondary antibodies at room temperature (RT) for 2 h. Secondary antibodies were Alexa Fluor 546-conjugated goat anti-rabbit IgG, goat anti-mouse IgG, goat anti-rat IgG (1:1,000; Invitrogen), Alexa Fluor 488-conjugated goat anti-rabbit IgG or goat anti-mouse IgG (1:300; Invitrogen), and Alexa Fluor 594-conjugated goat anti-rabbit IgG or goat anti-mouse IgG (1:1,000; Invitrogen). Nuclei were counterstained with DAPI (Sigma‒Aldrich), and cells were observed with a TE2000 Nikon microscope.

### 2.10 Immunohistochemistry of mouse brain tissue

The animals were anaesthetized with ketamine/xylazine (100 mg/kg and 10 mg/kg, respectively) and then transcardially perfused with saline. Brains were removed and fixed with 4% paraformaldehyde at 4 °C for 24 h and then kept in PBS at 4 °C. Forty-micron-thick sagittal sections were obtained using a vibratome (Leica) and subjected to immunohistochemistry. The brain sections were permeabilized with 1% Triton X-100 for 30 min at 25 °C. Nonspecific sites were blocked with 3% serum bovine albumin (Sigma‒ Aldrich) and 5% normal goat serum (Sigma‒Aldrich) diluted in PBS for 1 h before immunoreaction with the following antibodies: mouse anti-GFAP (1:1,000; Millipore, Cat. MAB360), rabbit anti-GFAP (1:1,000; DakoCytomation, Cat. Z0334), rabbit anti-Homer1 (1:500; Abcam, Cat. ab184955), synaptophysin (1:1000; Millipore, Cat. MABN1193) mouse anti-EGR1/Zif268 (1:200; Santa Cruz Biotechnology, Cat. Sc101033), rat anti-C3 (1:300, Abcam, Cat. ab11862), rabbit anti-S100a10 (1:1,000, Abcam, Cat. ab232524) at 4°C for 24 h. The sections were then washed with PBS and incubated with secondary antibodies at RT for 2 h. The secondary antibodies were Alexa Fluor 594-conjugated goat anti-rabbit IgG, Alexa Fluor 594-conjugated goat anti-mouse IgG, (1:1,000; Molecular Probes), and Alexa Fluor 488-conjugated goat anti-rabbit IgG (1:300; Molecular Probes). The nuclei were counterstained with DAPI. The sections were mounted with mounting medium (DakoCytomation) and imaged under a confocal microscope (Leica TCS SPE).

### 2.11 AβO intracerebroventricular (i.c.v.) injections and drug treatment

Three-month-old Swiss mice were kept in groups of five animals per cage in standard housing conditions (12 h light/dark cycle with controlled room temperature and humidity) with ad libitum access to food and water. All mouse experiments were performed in accordance with the Principles of Laboratory Animal Care from the National Institutes of Health and were approved by the Institutional Animal Care and Use Committee of the Federal University of Rio de Janeiro. I.c.v. injections were performed as previously described [33]. Briefly, mice were anaesthetized using isoflurane 2.5% (Cristália, São Paulo, Brazil) in a vaporizer system (Norwell, MA) and were gently restrained during the i.c.v. procedure. Mice received i.c.v. injections of 10 pmol AβO or vehicle (final volume, 3 µL) through a 2.5 mm-long needle. The needle was inserted unilaterally 1 mm to the right of the midline point equidistant from each eye and 1 mm posterior to a line drawn through the anterior base of the eye, as previously described [33]. LASSBio-1911 was dissolved in dimethyl sulfoxide (DMSO) and diluted with a mixture of saline and PEG-400 (40%); mice received daily treatment with vehicle or LASSBio-1911 (50 mg/kg) by the intraperitoneal route for ten days prior to biochemistry and toxicological analysis. In another experimental set, animals received AβO i.c.v. infusion one hour prior to vehicle or LASSBio-1911 (i.p.) treatment for 7-10 days followed by behavioural and molecular tests.

### 2.12 Toxicological analysis

The body weights of mice treated with LASSBio-1911 (50 mg/kg) or vehicle by intraperitoneal (i.p.) route were assessed daily for 10 days. Parameters indicative of toxicity, such as abdominal constriction, ptosis, piloerection, tremors, paralysis, tremor, reduction of body tonus, convulsions (seizures), corneal reflex, touch response, tail grip, and mortality, were randomly observed during animal manipulations [36]. On the 8th day of observation, animals were anaesthetized and sacrificed by cervical displacement and subjected to laparotomy, which allowed blood collection and macroscopic observation (general aspect, colour and weight) of organs and/or glands, such as the heart, lung, liver, kidney, spleen and pancreas. The levels of urea, creatinine, aspartate aminotransferase (AST), alanine aminotransferase (ALT), cholesterol, triglycerides and low-density lipoprotein (LDL) were determined. All biochemical analyses were validated and performed via spectrophotometry in a semiautomatic biochemical analyser (Bioplus-2000) using commercial kits (Vida Biotecnologia Ltda. - BRASIL).

### 2.13 Novel object recognition test

The behavioural analyses were performed according to a previously described protocol [33]. Mice were initially placed in an open field arena and subjected to a 5-min habituation session. The number of crossings and rearings was determined. Mice were then exposed to a training phase in which animals explored two identical objects for 5 min, and the time spent exploring each object was determined. One hour after the training session, mice were again placed in the arena for the test session, and one of the two familiar objects used in the training session was replaced by a novel one. Again, the time spent exploring familiar and novel objects was determined, and all data were analysed. The arena and objects were thoroughly cleaned between trials with 40% ethanol to eliminate olfactory cues. The results were expressed as the percentage of time exploring each object during the training or test session and were analysed using a one-sample Student’s t test comparing the mean exploration time for each object with the fixed value of 50%. Animals that recognize the familiar object as such (i.e., learn the task) spend more time exploring the novel object.

### 2.14 Passive avoidance behaviour

The step-down apparatus consisted of a barred floor box (40 × 25 × 35 cm) with a platform (9.5 × 7 × 3 cm) in the centre. During the training session, the animals were placed on the platform, and the time to descend from the platform was recorded. As soon as the animals placed their four paws on the metal grid, an electric shock of 0.5 mA was administered for 2 s. The animals were then placed on the platform. The test session took place 24 h after training, when the animals were again placed on the platform and the latency to descend from the platform was measured. Data are presented as retention values (descent latency in the training session subtracted from latency in the test session for each animal).

### 2.15 Flow cytometry

Cell dissociation for flow cytometry was adapted from a previously described protocol [37]. Mice were anaesthetized with ketamine/xylazine (100 mg/kg and 10 mg/kg, respectively) and then transcardially perfused with saline. Brains were removed, dissected, and rinsed in PBS. After removing the meninges, the hippocampus was cut into small pieces using a sterile scalpel and centrifuged at 300 ×g for 2 min at RT, and the supernatant was carefully removed. Enzymatic cell dissociation was performed using 20 U/ml papain in Gey’s buffer for 20 min at 37 °C under agitation. Papain activity was stopped with 10% FBS in Gey’s buffer at 4 °C. Tissue was mechanically dissociated with the aid of a pipette and filtered (70 μM nylon mesh strainer) to remove tissue pieces. Cell suspensions were centrifuged at 300 ×g for 10 min at 4 °C and fixed with 2% paraformaldehyde at 4 °C for 2 h. Suspensions were washed 3 times with PBS followed by centrifugation and then kept in PBS at 4 °C. Cell suspensions were centrifuged and permeabilized with saponin (0.1% in PBS) for 15 min at RT, centrifuged and washed 3 times with PBS and incubated with glycine (1.5 mg/ml) for 15 min at RT. Cells were washed 3 times with PBS and incubated with blocking solution (PBS/BSA 1%) for 1 h at 37 °C and then with the following primary antibodies: mouse anti-GFAP (1:500; Millipore, Cat. MAB360), rabbit anti-C3 (1:25; Invitrogen, Cat. MA1-40046), rabbit anti-S100a10 (1:20; Abcam, Cat. ab232524) at 4 °C for 24 h. The cell suspension was centrifuged at 300×g for 5 min at 4 °C, washed 3 times with PBS and incubated with the following secondary antibodies at RT for 2 h: Alexa Fluor 488-conjugated goat anti-mouse IgG (1:200; Molecular Probes) and Alexa Fluor 633-conjugated goat anti-rabbit IgG (1:1,000; Molecular Probes). After 2 hours, the cell suspension was centrifuged and washed as previously described, and the cells were resuspended in 500 µl of PBS. Background staining for antibodies was determined in antibody nonconjugated cells and fluorochrome-conjugated isotype control cells. Cells were analysed on a FACS Canto II flow cytometry system (BD Biosciences). Data were analysed with FlowJo vX software.

### 2.16 Quantitative RT‒PCR (qPCR)

The cortical astrocytes were lysed with TRIzol® (Invitrogen), and total RNA was isolated and purified with Direct-zol™ MiniPrep Plus (Zymo Research, Irvine, CA, USA) according to the manufacturer’s protocol. The RNA was quantified using a NanoDrop ND-1000 spectrophotometer (Thermo Fisher Scientific, Waltham, MA, USA). Total RNA (1-2 μg) was reverse transcribed with a GoScript^TM^ Reverse Transcriptase cDNA reverse transcription kit according to the manufacturer’s instructions (Promega Corporation, an affiliate of Promega Biotecnologia do Brasil, Ltda). Primers were designed and synthesized by IDT-DNA (San Diego, CA, USA). The specific forwards and reverse oligonucleotides were as follows: S100a10 (F) GCT TAC GTT TCA CAG GTT TG; (R) CCT GCC ACT AGT GAT AGA AAG and GAPDH (F) AAG AAG GTG GTG AAG CAG GCA TCT, (R) ACC CTG TTG CTG TAG CCG TAT TCA. Quantitative real-time PCR was performed using Fast SYBR Green Master Mix qPCR Master Mix (Applied Biosystem^TM^); the cycling conditions were 95 °C for 20 sec and 40 cycles of 95 °C for 1 sec and 60 °C for 20 sec in the QuantStudio 7 Flex System (Applied Biosystem^TM^). The relative expression levels of the genes were calculated using the 2^−ΔΔCT^ method [38].

### 2.17 Measurement of HDAC activity

The assay was performed according to the manufacturer’s recommendations (Histone Deacetylase Assay kit, Sigma‒Aldrich, Cat. CS1010) using cell culture extract (*in vitro* analysis) or hippocampal extract (*in vivo* analysis).

### 2.18 Mouse cytokine proteome profiler

The cytokine profile was analysed according to the manufacturer’s instructions (Mouse Cytokine Array Panel A Kit, R&D Systems, Catalogue #: ARY006). In these assays, 300 μg of hippocampal protein extract from animals in each experimental group was used. The samples were combined with a cocktail of biotinylated detection antibodies, followed by incubation with a membrane containing capture antibodies that are specific to the target proteins. The bound proteins were then detected using chemiluminescent reagents, and the signal strength was directly proportional to the amount of analyte bound. The inflammatory factors that appeared to be labelled in all membranes used for the assay were used to generate the heatmap. Analysis was performed only on cytokines that produced a signal in both experiments.

### 2.19 Western blotting

The cell extract protein concentration was measured by a BCA Protein Assay Kit (Cole-Parmer). Forty micrograms protein/lane was electrophoretically separated on a 10% SDS polyacrylamide gel and electrically transferred onto a Hybond-P PVDF transfer membrane (Millipore) for 1.5 h. Membranes were blocked in PBS-milk 5% at RT for 1 h. Next, membranes were incubated in block solution overnight with the following primary antibodies: mouse anti-Histone H3 (1:200; Cell Signalling, Cat. #14269), rabbit anti-acetyl histone H3 (1:1,500; Cell Signalling, Cat. #9677), rabbit anti-GFAP (1:1000; DAKO Cytomation, Cat. Z0334), mouse anti-Synaptophysin (1:1,000; Millipore, Cat. MABN1193), rabbit anti-PSD-95 (1:1,000; Abcam, Cat. ab18258), rabbit anti-P65 (1:1,000; Abcam, Cat. ab16502) and rabbit anti-β-actin (1:5,000; Abcam, Cat. ab8227). Membranes were incubated for 1 h with IRDye 680CW goat anti-mouse antibody and IRDye 800CW goat anti-rabbit antibody (LI-COR, 1:15,000) and then scanned and analysed using Un-Scan-It gel version 6.1 (Silk Scientific).

### 2.20 Densitometric analysis of immunofluorescence

Densitometries for the immunohistochemistry and immunocytochemistry images were performed using integrated density values generated with the ImageJ program (National Institutes of Health, USA, RRID: SCR_003070) or through analysis of the intensity in Leica Application Suite X software (RRID: SCR_013673). Immunocytochemistry data were collected from at least 10 fields per coverslip. The integrated density value was divided by the number of cells in each field. Analyses were performed in duplicate, and the graphs represent the average from at least 3 independent experiments. Immunohistochemistry analyses were performed with at least 2-3 sections from each animal, and 3 images per section at 40X or 63X magnification in the striatum were analysed.

### 2.21 Synaptic puncta analysis

***In vitro*** synapse analysis was performed as previously described [39]. Briefly, neurons were randomly identified and selected if nuclei staining (DAPI staining) was at least two diameters away from the neighbouring neuronal nucleus. Neuronal cultures were analysed by immunostaining for the pre- and postsynaptic markers synaptophysin and spinophilin, respectively. The green and red channels were merged and quantified using the Puncta Analyser plugin from ImageJ software (NIH, USA). Approximately 10-35 images were analysed per experimental condition, and the exact number of neuronal cultures per experimental group is indicated in the figure legends.

***In vivo*** synapse analysis was performed using two to three images of at least 2 sections (of approximately 40 μm each) derived from the hippocampal CA1 region of each animal. *In vivo* quantification of excitatory synaptic density was analysed by immunostaining for the pre- and postsynaptic markers synaptophysin and homer, respectively. Analysis of the colocalization rate of those markers was measured by Leica Application Suite X software (RRID:SCR_013673).

### 2.22 Statistical analysis

GraphPad software, version 8.0 (GraphPad Software, La Jolla, CA, RRID:SCR_002798), was used for statistical analysis. Because all statistical tests involved multiple conditions, analysis of variance was applied in all comparisons, followed by Tukey’s posttest when statistical significance was achieved. A confidence interval of 95% was used, and a *p* value < 0.05 was considered statistically significant. Densitometry of blotted gels was performed using Un-Scan-It gel version 6.1 (Silk Scientific, Inc., Orem, UT). Data are reported as the mean ± SEM, and error bars in the graphs represent the SEM.

## 3. RESULTS

### 3.1 LASSBio-1911 modulates astrocyte reactivity and improves synaptic density

LASSBio-1911 (**Figure 1A**) is a novel *N*-acylhydrazone derivative previously designed from isosteric changes in the structure of the natural product trichostatin A (TSA), a well-known nonselective HDAC inhibitor [40]. However, unlike TSA, LASSBio-1911 was characterized as a highly potent inhibitor of HDAC6 (IC_50_ = 15 nM) with 15-fold selectivity over HDAC8 (IC_50_ = 230 nM) and no significant activity on other class I HDAC isoforms (Rodrigues et al. 2016). Molecular docking studies allowed us to identify the main interactions involved in the molecular recognition of LASSBio-1911 by HDAC6, which consisted of a bidentate coordination of the hydroxamate group with the zinc ion, stabilized by additional hydrogen bond interactions with His-573, His-574 and Tyr-745 (**Figure 1B-C).** In addition, π-π stacking interactions of the phenyl group attached to the hydroxamate of LASSBio-1911 with Phe-583 and Phe-643 and a hydrogen bond interaction of the N-H group of the *N*-acylhydrazone moiety with Ser-531 in the entrance of the active site were observed (**Figure 1B-C**).

**Figure 1:**
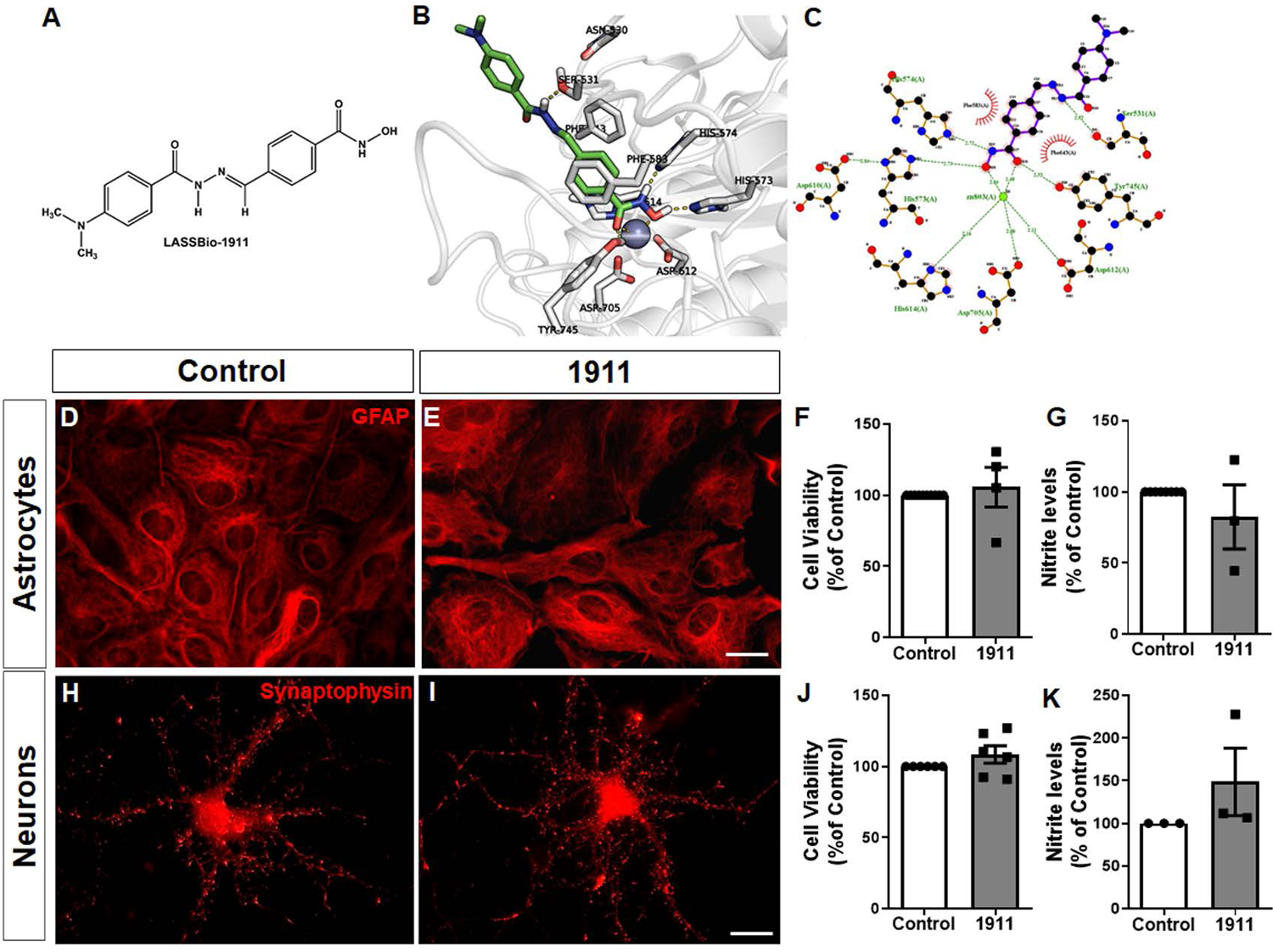
LASSBio-1911 is not toxic to neural cells in vitro. Chemical structure of N-acylhydrazone derivative LASSBio-1911 **(A)** and interaction mode of LASSBio-1911 (carbons in green) in HDAC6 (carbons in white) **(B)**. The three-dimensional interaction mode of LASSBio-1911 can be visualized in **B** (generated by PyMol v. 1.5.0.5) and its schematic representation in **C** (generated by LigPlot^+^ [79, 80]. **D, E, H, I:** Representative images of astrocytes and neuronal cultures treated with 1 μM LASSBio-1911 **(1911)** and vehicle **(Control)** for 24 hours and stained with anti-GFAP **(D, E)** and anti-synaptophysin **(H, I)**. Toxicity was evaluated by measurement of cell viability (MTT assays) and nitrite levels**. F, G:** Quantification of cell viability and nitrite levels in astrocyte cultures (n=4); **J, K:** Quantification of cell viability and nitrite levels in neuronal cultures (n=4). Student’s *t* test. analysis. Scale bars: 20 μm **(E)**; 20 μm **(I)**.

To address LASSBio-1911 toxicity, we first evaluated its impact on the viability of astrocytes and neuronal cultures derived from the murine hippocampus (**Figure 1D-K**). Cultures were treated with 1 μM LASSBio-1911 for 24 hours followed by assessment of MTT assays and nitrite production. The concentration of LASSBio-1911 used in this work was determined by the maximum concentration that had no impact on astrocyte viability. Treatment of astrocytes with LASSBio-1911 did not impact the morphology of astrocytes, which presented a protoplasmic morphology with GFAP-labelled filaments throughout the cytoplasm, characteristic of those cells (**Figure 1D, E)**. Similarly, LASSBio-1911 induced no change in neuronal morphology, as exhibited by the presence of long synaptophysin-labelled processes (**Figure 1H, I)**. Additionally, adult Swiss mice treated with LASSBio-1911 (50 mg/kg, i.p., for one week) showed no changes in body weight, relative weight of several vital organs or in biochemical markers of renal and hepatic function. Altogether, these data showed that the novel HDAC inhibitor LASSBio-1911 presented no toxicity in the *in vitro* and *in vivo* assays performed in this work.

Phenotypic heterogeneity of astrocytes is mediated by different pathological stimuli [15, 41]. To evaluate the impact of LASSBio-1911 on the astrocytic phenotype, we used flow cytometry and immunohistochemistry of hippocampal cell suspensions and tissues, respectively. (Figure 2). Using suspensions of hippocampal cells from LASSBio-1911-treated mice, we found a decreased number of GFAP/C3-positive cells (**Figure 2D-F),** a neurotoxic astrocytic profile, and an increased number of GFAP/S100a10-positive cells (**Figure 2G-I**), a neuroprotective astrocytic phenotype. Corroborating these data, we also found that the hippocampus of animals treated with LASSBio-1911 showed decreased colocalization between GFAP/C3 and increased colocalization between GFAP/S100a10 (**Figure 2A-C**).

**Figure 2:**
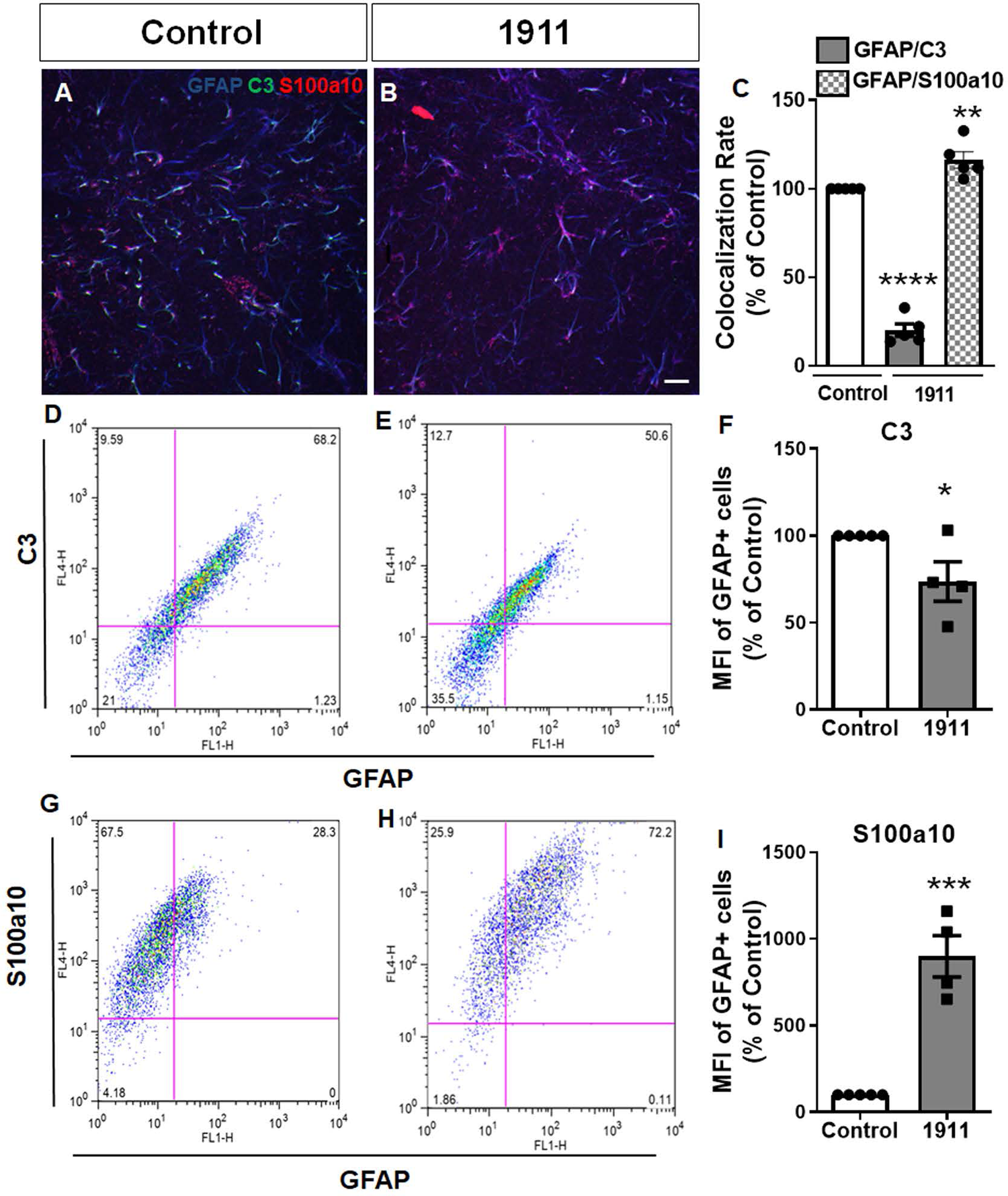
LASSBio-1911 decreases astrocyte reactivity markers in mice. **A, B:** Representative images of GFAP, C3 and S100a10 immunostaining hippocampal sections of Swiss adult mice treated with vehicle **(Control)** or LASSBio-1911 **(1911). C**: Quantification of the colocalization rate of C3/GFAP and S100a10/GFAP in the hippocampus of different groups (n=5 animals per group); **D-I:** FACS profile quantification of GFAP/C3 **(D-F)** and GFAP/S100a10 **(G-I)** cells isolated from hippocampal tissues from vehicle- and LASSBio-1911-treated mice (n=4 animals per group). Student’s *t* test, *P < 0.050; ****P < 0.0001; ***P < 0.001. Scale bar: 20 μm **(B)**.

Synapse damage is an important feature of memory-related diseases [10]. To assess the effect of LASSBio-1911 on neuronal synapse density, we evaluated the levels and distribution of pre(synaptophysin) and post (spinophilin, PSD-95 or Homer-1) synaptic proteins in hippocampal neuronal cultures (**Figure 3A-F**) and in the CA1 region of the mouse hippocampus (**Figure 3G-L**). LASSBio-1911 increased the number of colocalized synaptic puncta by 30 and 77% *in vitro* (**Figure 3A-C**) and *in vivo* (**Figure 3G-I**), respectively. Western blotting assays of the hippocampal tissue of LASSBio-1911-injected mice also showed increased levels of the presynaptic protein synaptophysin (**Figure 3L**).

**Figure 3:**
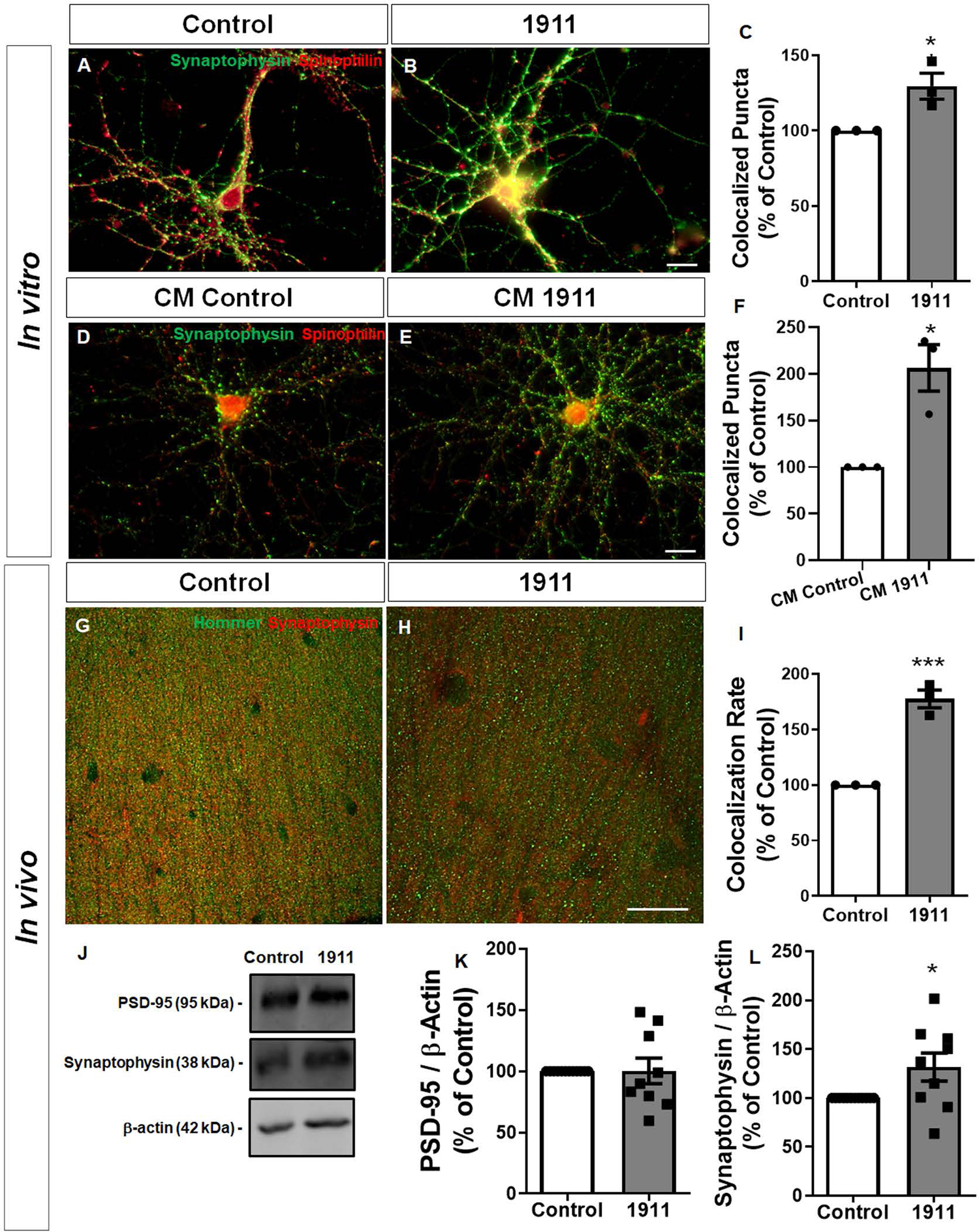
LASSBio-1911 increases synaptic protein levels and distribution in cultured neurons and in mice. **A, B:** Representative images of synaptophysin/spinophilin immunostaining of hippocampal neuronal cultures treated with vehicle **(Control)** or LASSBio-1911 **(1911). C**: Quantification of the colocalization rate of synaptophysin/spinophilin puncta (n=3 cultures); **D, E:** Representative images of synaptophysin/spinophilin immunostaining of hippocampal neuronal cultures treated with conditioned medium from nontreated astrocytes **(CM Control)** or from astrocytes treated with LASSBio-1911 **(CM 1911); F**: Quantification of the colocalization rate of synaptophysin/spinophilin puncta (n=3 cultures); **G-H:** Representative images of synaptophysin/hommer immunostaining of hippocampal sections of Swiss adult mice treated with vehicle **(Control)** or LASSBio-1911 **(1911). I:** Quantification of the colocalization rate of synaptophysin/hommer puncta (n=3 animals per group). **J:** Representative images and quantification of Western blotting gels of hippocampal tissues isolated from Swiss adult mice injected with vehicle **(Control)** or LASSBio-1911 **(1911)** and stained with anti-PSD-95 **(K)** and anti-synaptophysin **(L)** (n=9 animals per group). β-Actin was used as a loading control for Western blotting. LASSBio-1911 increased synaptic protein colocalization puncta in mice. Student’s *t* test, *P < 0.050; ***P < 0.001. Scale bars: 20 μm **(B)**; 20 μm **(E);** 20 μm **(H)**.

To determine whether LASSBio-1911 modulates the synaptogenic potential of astrocytes, purified neuronal cultures were treated with conditioned medium from control astrocytes (CM Control) or with conditioned medium from astrocytes treated with LASSBio-1911 (CM 1911), and synapse density was further evaluated. We found that compared to nontreated astrocytes, astrocytes treated with LASSBio-1911 exhibited a twofold increase in synaptogenic potential (**Figure 3D-F**).

Altogether, these data suggest that LASSBio-1911 has neuroprotective potential since it modulates astrocyte reactivity, enhances astrocytic synaptogenic properties, and improves synaptic density both in cultured neural cells and in the mouse brain.

### 3.2 LASSBio-1911 modulates histone acetylation in healthy and AβO-exposed mouse brains

The HDAC inhibitory activity of LASSBio-1911 was previously determined by Eurofins Discovery (https://www.eurofinsdiscovery.com/) using recombinant human HDAC isoforms [20]. To characterize HDAC activity in brain tissue, we measured HDAC activity in hippocampal extracts from vehicle- or LASSBio-1911-treated mice (**Figure 4A**). We found that hippocampal tissue of mice treated with LASSBio-1911 showed a 32% decrease in HDAC activity (**Figure 4B**) and a 58.5% increase in the levels of H3 acetylated histones (**Figure 4C**). These data support the previous demonstration that LASSBio-1911 is an HDAC inhibitor and demonstrate for the first time its effect on brain HDACs.

**Figure 4:**
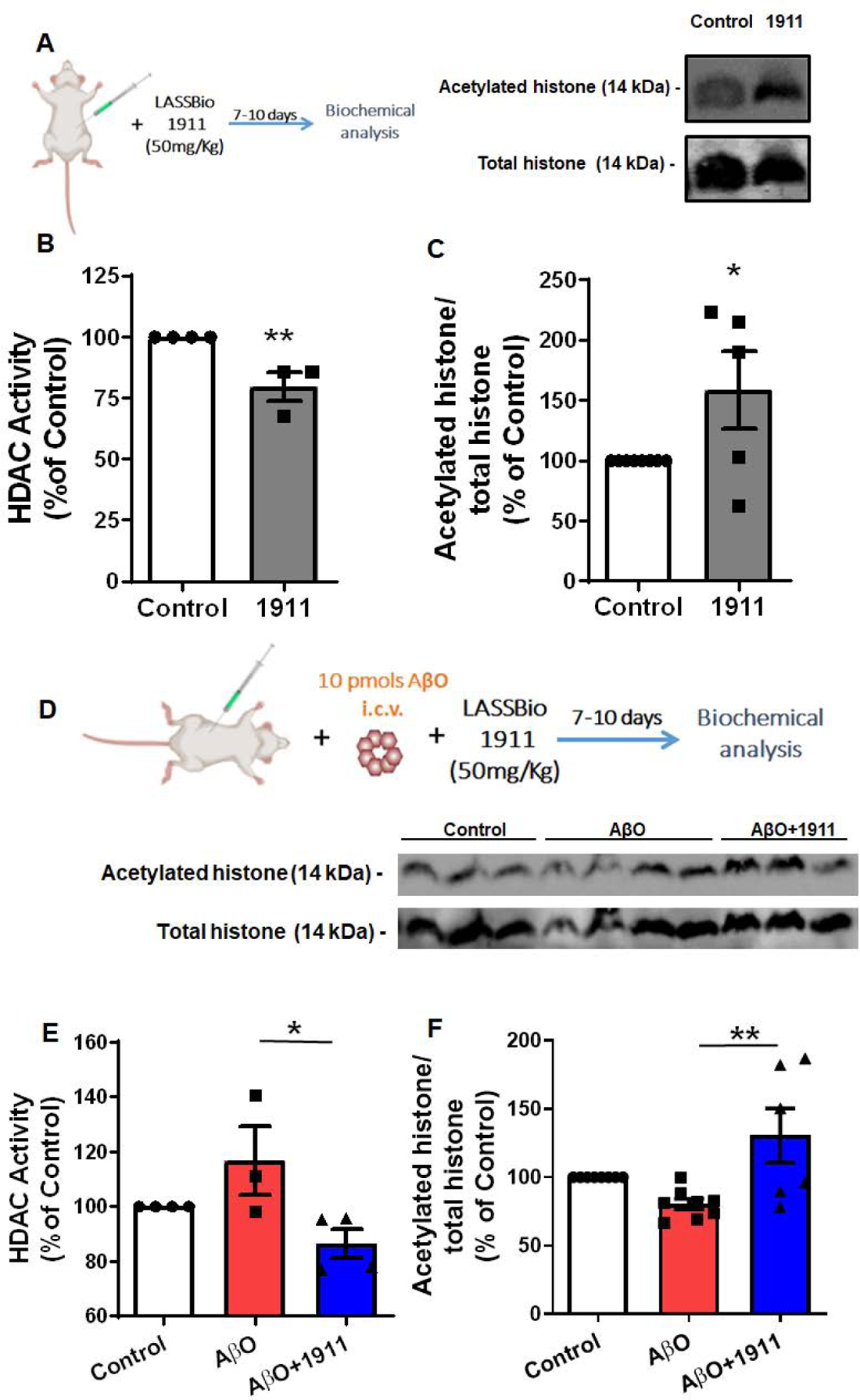
LASSBio-1911 rescues the effect of A βO on HDAC activity and histone acetylation in the brain. **A:** Schematic of LASSBio-1911 treatment of Swiss adult mice; **B:** Quantification of HDAC activity of hippocampal tissues derived from Swiss adult mice injected with vehicle **(Control)** or LASSBio-1911 **(1911)** (n=4 animals per group); **C:** Representative images and quantification of Western blotting gels of hippocampal tissues isolated from Swiss adult mice injected with vehicle **(Control)** or LASSBio-1911 **(1911)** and stained with anti-acetylated histone. Total histones were used as a loading control for Western blotting. The ratio of acetylated histones to total histones was used to evaluate the level of acetylated histones (n=5 animals per group). Student’s *t* test, *P < 0.05; **P < 0.01. **D:** Schematic of AβO injection and LASSBio-1911 treatment of Swiss adult mice; **E:** Quantification of HDAC activity of hippocampal tissues derived from Swiss adult mice injected with vehicle **(Control),** AβO **(A**β**O)** or LASSBio-1911 and AβO **(A**β**O +1911)** (n=4 animals per group); **F:** Representative images and quantification of Western blotting gels of hippocampal tissues isolated from Swiss adult mice injected with vehicle **(Control),** AβO **(A**β**O)** or LASSBio-1911 and AβO **(A**β**O+1911)** and stained with anti-acetylated histone. Total histone was used as a loading control for Western blotting. The ratio of acetylated histones to total histones was used to evaluate the level of acetylated histones (n=6 animals per group). Comparisons between multiple groups were analysed using one-way ANOVA followed by Tukey’s *post hoc* tests *P < 0.05; **P < 0.01.

Inhibitors of HDAC have emerged as promising targets for the treatment of neurodegenerative diseases [42]. In addition, evidence has demonstrated that reduced levels of HDAC6 in the mouse brain resulted in memory and learning restoration in a mouse model of AD [43]. Considering this finding, we evaluated the efficacy of LASSBio-1911 in modulating acetylation in AD using an experimental model of AβO toxicity [14, 33]. AβO accumulate in the brains of AD patients and are increasingly considered major toxins leading to AD-associated neuronal dysfunction.

We found that i.c.v. infusion of 10 pmol AβO in mice increased HDAC activity in the hippocampal tissue by approximately 20%, whereas systemic treatment with LASSBio-1911 resulted in a decrease in enzyme activity (**Figure 4E**). Similarly, Western blotting analyses further showed that LASSBio-1911 treatment not only restored but also improved the hippocampal levels of H3 acetylated histone (**Figure 4F**). Altogether, these results show that LASSBio-1911 controls acetylation levels in the brain and mitigates AβO-induced effects in the hippocampus.

### 3.3 LASSBio-1911 mitigates the astrocytic reactivity profile triggered by AβO in mice

Astrocyte dysfunction and reactivity have been considered important players in AD. Efforts to manipulate the astrocytic reactivity phenotype have emerged as a potential alternative to AD therapy [19, 44]. Here, we found that exposure of astrocytes to AβO led to a 36% decrease in the levels of acetylated histones, suggesting that epigenetic control in astrocytes may play an important role in AD and thus be a target for LASSBio-1911.

We then asked whether LASSBio-1911 would rescue the effects of AβO on astrocytes. We previously demonstrated that AβO induce morphological and functional alterations in astrocytes, such as astrocyte process atrophy and decreased production of synaptogenic cytokines, leading to decreased synaptogenesis properties of those cells (Diniz et al. 2017). Here, we extended this evaluation by showing that hippocampal tissue from AβO-infused mice exhibited a range of modifications in the level of several cytokines, such as ILs (IL-1α, IL-1β, IL-13), TNF-α and others. Interestingly, LASSBio-1911 systemic treatment rescued the cytokine profile induced by AβO in mice (**Figure 5A-C**).

**Figure 5:**
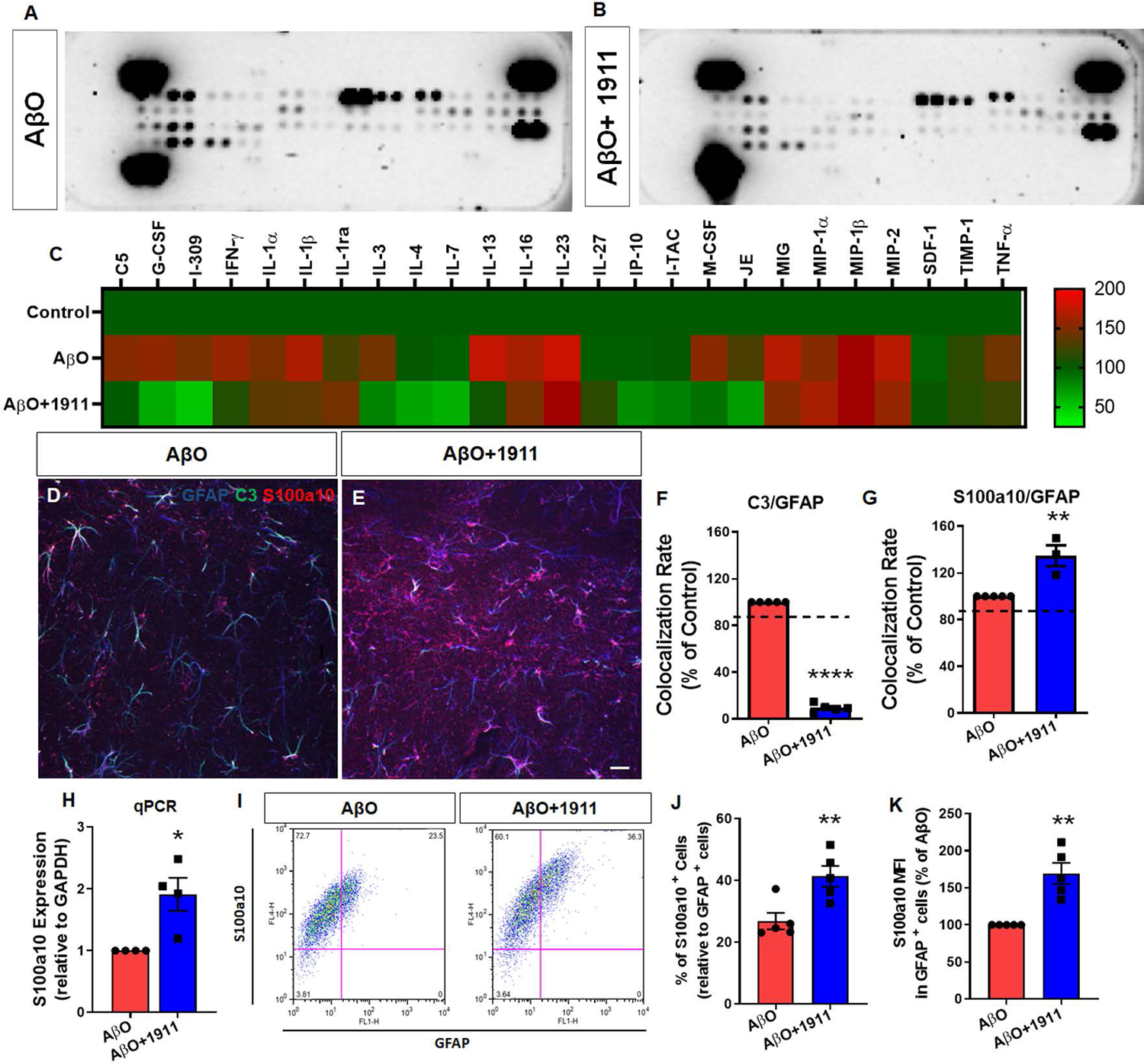
LASSBio-1911 rescues the astrocytic reactivity profile triggered by AβO in mice. **A-C:** Proteomic profile of hippocampal tissues of Swiss adult mice injected with vehicle **(Control),** AβO (AβO) or LASSBio-1911 and AβO (AβO**+1911); D, E:** Representative images of GFAP, C3 and S100a10 immunostaining hippocampal sections of Swiss adult mice treated with AβO and AβO + 1911**; F, G:** Quantification of the colocalization rate of C3/GFAP **(F)** and S100a10/GFAP **(G)** in the hippocampus of different groups (n=5 animals per group). The dashed lines refer to control values. **H:** qPCR assays of the levels of S100a10 in the hippocampus of AβO- and AβO+LASSBio-1911-injected mice (n=4 animals per group); **I-K:** FACS profile quantification of S100a10/GFAP cells isolated from hippocampal tissues from AβO- and AβO+LASSBio-1911-injected mice (n=5 animals per group). K shows the intensity of S100a10 staining per GFPA+ cell. Student’s *t* test *P < 0.05; **P < 0.01; ****P < 0.0001 Scale bar: 20 μm **(E)**.

To correlate such profiles with the astrocyte reactivity phenotype, we addressed the levels of the astrocytic neurotoxic marker C3 and the astrocytic neuroprotector marker S100a10 in the hippocampus of AβO-injected and LASSBio-1911-treated mice. LASSBio-1911 reduced the colocalization rate of C3/GFAP-positive cells by 90% in the hippocampus of AβO-injected mice (**Figure 5D, E, F)** but increased the colocalization rate of S100a10/GFAP cells by 34% (**Figure 5D, E, G)**. The upregulated expression of S100a10 in the hippocampus of the animals was also confirmed by qPCR (**Figure 5H**). Flow cytometry assays revealed that not only the number of S100a10-GFAP cells but also the intensity of S100a10 per cell was increased by LASSBio-1911 (**Figure 5I-K**).

To correlate the effects of LASSBio-1911 on astrocytes on synaptic deficits induced by AβO, cultured hippocampal astrocytes were treated with vehicle or LASSBio-1911, and the conditioned media from these cultures were collected. We thus evaluated the ability of these CMs to protect neurons against AβO cytotoxicity. As observed in **Figure 6**, the conditioned medium from astrocytes treated with LASSBio-1911 was more efficient in protecting neurons against AβO synaptic toxicity (**Figure 6B-E**). Altogether, these data show that AβO triggers an astrocytic reactivity phenotype profile in mice that can be mitigated by LASSBio-1911.

**Figure 6:**
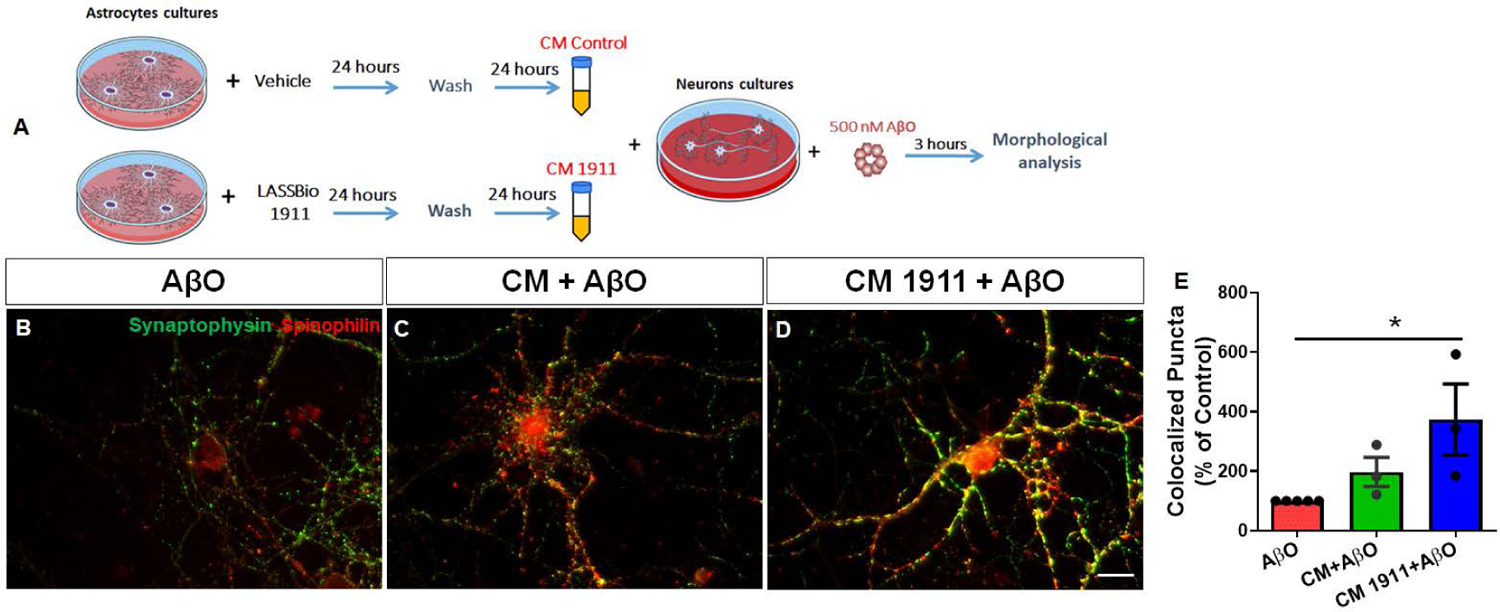
LASSBio-1911 enhances the neuroprotective potential of astrocytes against AβO. **A:** Schematic of astrocyte conditioned media preparation and neuroprotection assay against AβO. Purified neuron cultures were treated with AβO in the presence of serum-free medium, conditioned medium from control astrocytes **(CM Control),** and conditioned medium from astrocytes treated with LASSBio-1911 **(CM 1911), and** the density of glutamatergic synapses was determined by immunocytochemistry after 3 hours. **B-D:** Representative images of synaptophysin/spinophilin immunostaining of hippocampal neuronal cultures maintained in the presence of AβO and serum-free medium **(AβO),** conditioned medium from control astrocytes **(CM Control+AβO),** and conditioned medium from astrocytes treated with LASSBio-1911 **(CM 1911+AβO). E**: Quantification of the colocalization rate of synaptophysin/spinophilin puncta (n=4 cultures). Comparisons between multiple groups were analysed using one-way ANOVA followed by Tukey’s *post hoc* tests *P < 0.050. Scale bar: 20 μm.

### 3.4 LASSBio-1911 improves behavioural performance and mitigates synaptic and memory function loss in AβO-injected mice

Synapse and memory loss is well characterized in several animal models of AD [45]. We and others have previously shown that AβO injection elicits memory deficits in mice [14, 33]. We thus used the AβO toxicity animal model to evaluate whether LASSBio-1911 might prevent AD-associated cognitive dysfunction. Then, we carried out behavioural tests in vehicle- or LASSBio-1911-treated animals. Hippocampal-dependent perirhinal cortex-dependent recognition memory was assessed using the novel object recognition task (**Figure 7**).

**Figure 7.**
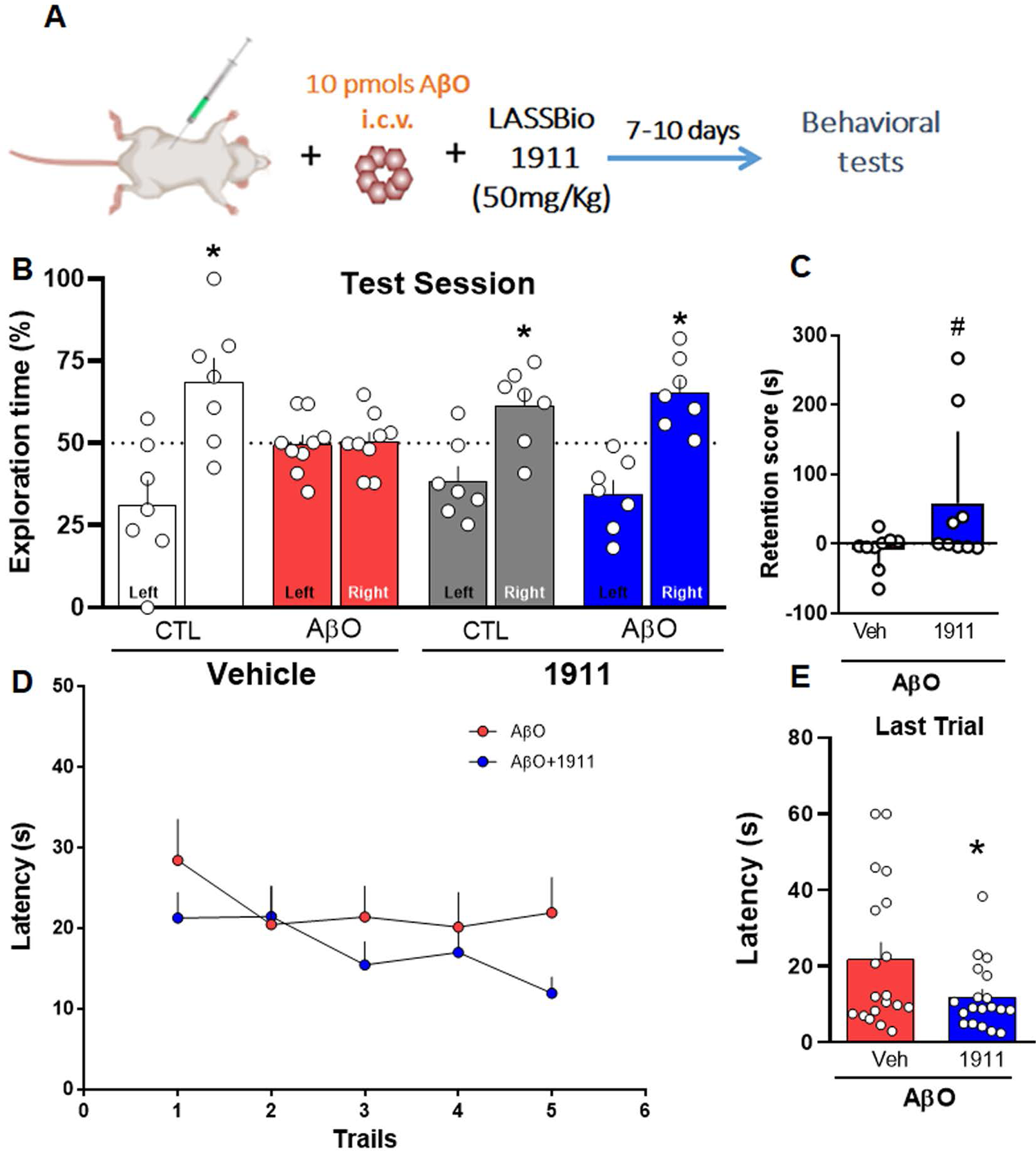
LASSBio-1911 rescues memory deficits induced by AβO. **A:** Schematic of AβO injection and LASSBio-1911 treatment of Swiss adult mice. Mice were infused with AβO (10 pmol/site) and treated with LASSBio-1911 (50 mg/kg, once a day) for 7-10 days. Forty-eight hours after LASSBio-1911 treatment, animals were exposed to three different tasks to evaluate cognitive function. **B:** Mice were tested in the NOR test. One-sample Student’s t test compared with the chance level of 50% (n=7-9 mice per group, *p < 0.05). **C:** Retention score (sec) of mice in the step-down inhibitory avoidance; Student’s t test; n = 9 mice per group). **D:** Escape latency across 5 consecutive training trials **(F)** and time spent in the target quadrant during the probe trial **(E)** of the MWM test. Repeated measures ANOVA followed by Tukey’s test (n=19 mice per group, *P < 0.05).

For these experiments, LASSBio-1911 was intraperitoneally administered 30 min before the infusion of 10 pmol AβO (i.c.v.). As expected, vehicle-infused mice learned the novel object recognition (NOR) memory task, as demonstrated by longer exploration of the novel object over the familiar object (**Figure 7B, white bars)**. In contrast, AβO-infused mice failed the NOR task (**Figure 7B, red bars)**. Remarkably, treatment with LASSBio-1911 mitigated AβO-induced cognitive impairment in mice (**Figure 7A-C**). Confirming this result, compared to control mice, mice infused with AβO (10 pmol/site, i.c.v.) showed a higher latency to find the submerged platform in sessions of MWM training (**Figure 7D**). Additionally, AβO-infused mice showed reduced memory retention, as indicated by the decreased time spent by these animals in the target quadrant during the probe trial compared to controls (**Figure 7E**). These results demonstrate that LASSBio-1911 improves the learning parameters and memory performance of AβO-injured mice.

Synapse loss is an important hallmark of several memory dysfunction diseases. We thus investigated whether the effect of LASSBio-1911 on memory recovery was associated with morphological synaptic changes. As expected, we first observed decreased colocalization between the presynaptic marker synaptophysin and the postsynaptic marker Homer-1 in the hippocampus of AβO-infused mice (**Figure 8B-D**). These data were corroborated by Western blotting assays that showed decreased levels of the post- and presynaptic proteins PSD-95 and synaptophysin, respectively, in hippocampal lysates of AβO-infused mice (**Figure 8H-J**). Interestingly, the reduction in the number of colocalized Homer-1/Synaptophysin synaptic puncta (**Figure 8D**) and in the levels of PSD-95 and synaptophysin (**Figure 8H-J**) induced by AβO was prevented by LASSBio-1911 systemic treatment.

**Figure 8:**
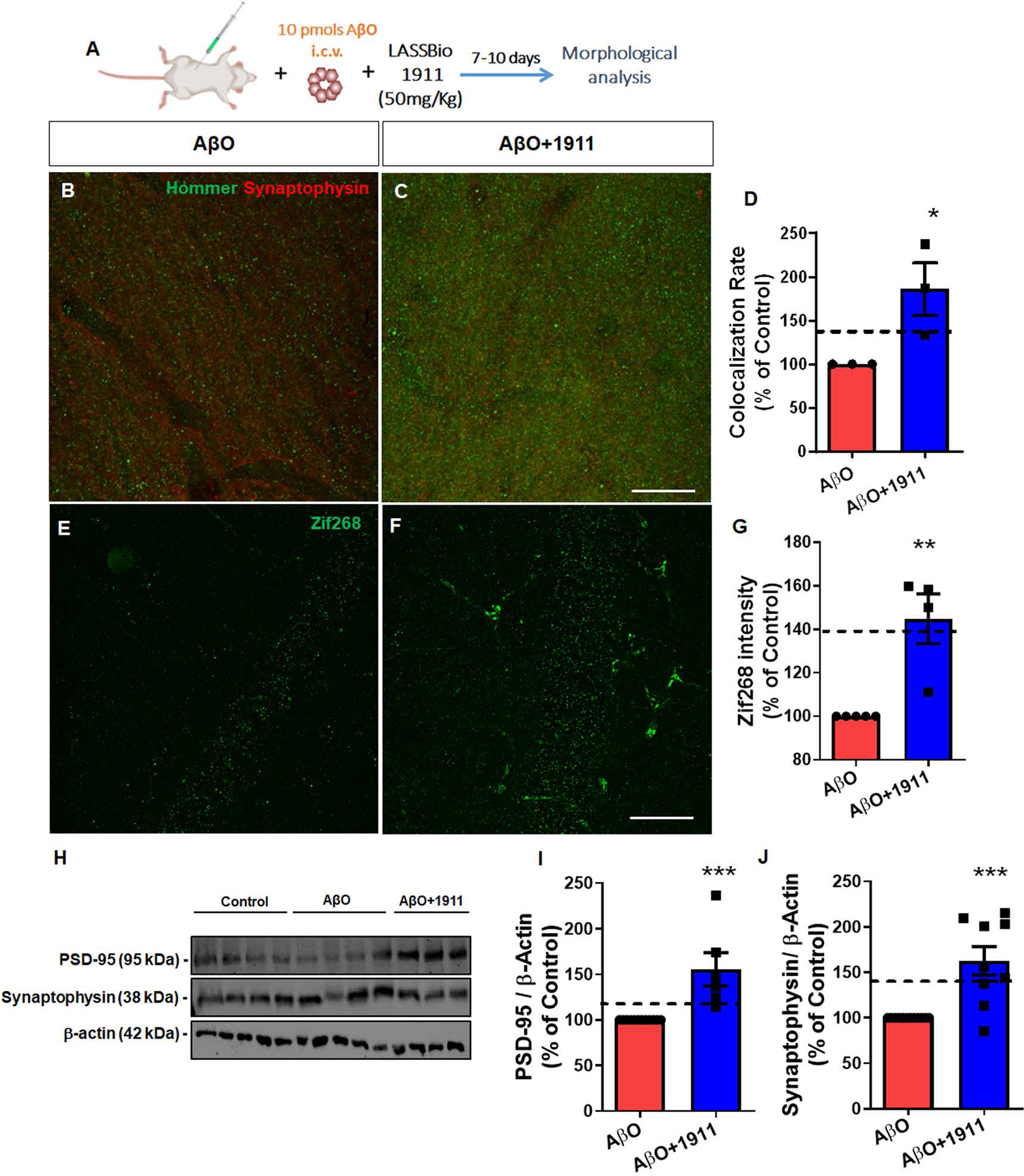
LASSBio-1911 rescues synaptic and memory deficits induced by AβO. **A:** Schematic of AβO injection and LASSBio-1911 treatment of Swiss adult mice; **B, C:** Representative images of synaptophysin/hommer immunostaining of hippocampal sections of Swiss adult mice treated with AβO **(AβO)** or LASSBio-1911 and AβO **(AβO+1911). D:** Quantification of the colocalization rate of synaptophysin/hommer puncta (n=3 animals per group). Dashed lines represent control values; **E, F:** Representative images of Zif268/EGR1 immunostaining of hippocampal sections of Swiss adult mice treated with AβO **(AβO)** or LASSBio-1911 and AβO **(AβO+1911). G:** Quantification of the Zif268/EGR1 intensity (n=5 animals per group); **H-J:** Representative images and quantification of Western blotting gels of hippocampal tissues isolated from Swiss adult mice injected with AβO **(AβO)** or LASSBio-1911 and AβO **(AβO+1911)** and staining for PSD-95 and synaptophysin (n=9 animals per group). Student’s *t* test *P < 0.05; **P < 0.01 ***P < 0.001. Scale bar: 20 μm.

To investigate the ability of LASSBio-1911 to modulate synaptic activity, we evaluated the distribution of early growth response transcription factor 1 (EGR1; also known as Zif268/EGFR1) in neurons of the molecular layer of the hippocampal region CA1 (**Figure 8E-G; Zif268)**. Zif268/EGR1 is associated with mechanisms underlying neurotransmission and neuronal plasticity, including those related to neuronal activity associated with learning and memory [46]. To do that, tissues were immunolabelled against the Zif268/EGR1 protein. It is worth mentioning that Zif268/EGFR1 staining was strongly observed in blood vessels (data not shown). The levels of Zif268/EGR1 staining were evaluated specifically in the molecular layer of CA1, where most of the hippocampal neuronal nuclei are located. As observed in **Figure 8E-G**, treatment with LASSBio-1911 elicited a 40% increase in Zif268/EGFR1 staining in the CA1 of the AβO-injected animals compared to animals in the control group. Altogether, these data demonstrated that LASSBio-1911 treatment improves behavioural performance and rescues synaptic loss induced by AβO in mice.

In summary, these data suggest that LASSBio-1911 mitigates synapse and memory loss induced by AβO and modulates the neuroprotective potential of astrocytes.

## 4. DISCUSSION

In this study, we provided evidence that LASSBio-1911, a novel HDAC6-selective inhibitor, modulates astrocyte reactivity, improves astrocyte synaptogenic properties and rescues synaptic loss and memory function in an animal model of Alzheimer’s disease. Together, the results presented here revealed a new HDAC inhibitor that mitigates AβO toxicity by modulating astrocyte phenotype and function.

Alzheimer’s disease is one of the biggest public health concerns worldwide. AD is a multifactorial disease that includes a series of cellular and tissue alterations, such as oxidative stress, synaptic dysfunction, glial reactivity, and neuroinflammation, which can result in neuronal death and memory loss [47]. Although numerous clinical and preclinical studies have been conducted to identify an effective treatment for AD, only a very limited number of drugs have been clinically applied thus far. In this scenario, drugs such as inhibitors of HDACs, with multiple intracellular targets, have emerged as alternative treatments for multifactorial diseases. However, the development and use of such drugs is still a major challenge due to their diverse responses and toxicity [48]. Although the cytotoxic effect of LASSBio-1911 has been previously shown for a prostate cancer cell line [49], we did not observe LASSBio-1911 toxicity in cultured primary neural hippocampal cells or in mice.

A decrease in the levels of histone acetylation [50] has been associated with decreased memory performance in aged mice, although there is still no consensus in the literature about the pattern of histone acetylation in AD models [51]. In the present work, we found that cultured astrocytes and animals i.c.v infused with AβO showed a decrease in the level of acetylated histones, indicating epigenetic dysfunction in this model. This finding corroborates previous studies that indicated a decrease in histone acetylation and gene expression levels in AD models [52–54]. Consistent with previous studies [55, 56], we also found an increase in the levels or overall activity of HDACs in *in vitro* and *in vivo* models of AβO toxicity. As expected, LASSBio-1911 decreased HDAC activity and increased the levels of histone acetylation *in vitro* and *in vivo*. These data support the previous demonstration that LASSBio-1911 is an HDAC6 inhibitor and demonstrate, for the first time, its effect on HDAC activity in the brain.

Inhibition of HDACs ameliorates synaptic loss and neuroinflammation and reverses cognitive deficits in AD models [51, 57–59], which broadens the impact of LASSBio-1911 in the brain. HDAC6 has been reported to be involved in many cellular events controlled by deacetylation of histone and nonhistone proteins, such as microtubule dynamics and axonal transport, apoptotic cell death, misfolded protein clearance, cellular stress prevention [60–62] and chaperone expression [63]. It remains to be investigated whether LASSBio-1911 exerts its action by affecting these events.

In the present study, we have shown that LASSBio-1911 regulates synaptic protein expression and synapse formation through the modulation of astrocytic phenotype and function. However, the specific role of HDACs in astrocytes, mainly in AD astrocytes, remains largely unexplored. Only in the last decade has it been found that astrocytes and microglia express HDACs [64–66]. Treatment with HDAC paninhibitors, such as SAHA and ITF2357, increases histone acetylation in glial cells, in addition to inhibiting LPS-induced expression of glial proinflammatory markers [67]. This evidence demonstrates that the modulation of nuclear histone acetylation levels in glial cells plays an important role in modulating inflammation. In agreement with this, we found that LASSBio-1911 reduced the levels of several inflammation markers in the hippocampus of AβO-injected mice, including TNF-α, IFN-γ, and many interleukins, suggesting that not only the global inhibition of HDAC has a beneficial effect on inflammation but also the selective inhibition of HDAC6.

In pathological models, astrocytes may show a molecular signature associated with altered morphology and physiological properties, a condition known as astrocytic reactivity [66]. Although those subpopulations of astrocytes may considerably vary, two of them have been recognized as A1 and A2 reactive astrocytes, which are characterized by the upregulated expression of C3 and S100a10, respectively. A1 astrocytes produce large amounts of inflammatory molecules, such as several cytokines and complement system cascade proteins, which have previously been shown to be destructive to synapses [15]. In contrast, A2 astrocytes are highly capable of releasing neurotrophic factors to support neuronal survival [68].

The modulation of astrocytic reactivity has led to contradictory results in the literature. Suppression of the JAK2-STAT3 pathway, which is required for the induction and maintenance of the reactive phenotype, ameliorated synaptic and memory deficits in mouse models of AD [44]. In contrast, the ablation of reactive astrocytes exacerbated disease pathology in a model of AD due to impaired amyloid clearance and increased neuroinflammation [69]. In AD specifically, studies have shown altered levels of synaptogenic molecules released by astrocytes, such as thrombospondin (TSP-1) [70] and TGF-β1 [14], and an increased proportion of A1 astrocytes, causing neuroinflammation and loss of synaptic properties [15].

Within this context, the search for new therapeutic strategies focused on the manipulation of astrocytic phenotypes may be conducted with two objectives: 1) to mitigate neuroinflammation and 2) to restore astrocytic synaptogenic and neuroprotective properties [71]. Here, we showed that while AβO induces a neurotoxic astrocytic profile, LASSBio-1911 rescues the neuroprotective phenotype and mitigates the levels of inflammation. Within a similar approach, glucagon-like peptide-1 receptor (GLP-R1) agonists have recently been shown to block microglia-mediated A1 astrocyte formation, prolong life and reduce behavioural deficits and neuropathological abnormalities in animal models of PD [72] and AD [73], indicating an effect of these drugs via microglia. Here, we found a reduction in the C3-positive astrocytic population and an increase in the S100a10-positive astrocytic population, suggesting a conversion from the A1 to A2 phenotype induced by LASSBio-1911. These results suggest that this compound can induce a shift in the astrocytic phenotype, which impacts the levels of inflammation. Although we cannot completely rule out an effect of LASSBio-1911 via microglia, our *in vitro* findings using microglia-free cultures indicate that the drug exerts a direct effect on astrocytes. Our data are in agreement with recent evidence that inhibition of HDAC6 blocks inflammatory signalling and decreases the levels of inflammatory cytokines, such as IL-1β and TNF-α, in other systems [74, 75]. Our work is, however, the first to show that inhibition of HDAC6 in the brain has a major therapeutic impact on inflammatory control in a neurodegenerative animal model.

Here, we verified that LASSBio-1911, in addition to rescuing the astrocytic reactivity phenotype, improves astrocytic synaptogenic and synaptoprotective properties. Several studies support the role of astrocytes as modulators of excitatory and inhibitory neural circuits in the brain [18, 34, 76–78]. In brain ageing and in AD in particular, studies have shown altered levels of synaptogenic molecules released by astrocytes, such as thrombospondin (TSP-1) and TGF-β1 (Jayakumar et al. 2014; Diniz et al. 2017; Matias et al. 2022). Histone deacetylase inhibitors have previously been shown to upregulate astrocytic GDNF and BDNF gene transcription and to protect dopaminergic neurons [25]. Although further studies are needed to identify the exact mechanisms by which LASSBio-1911 enhances synapse formation, the fact that conditioned medium derived from LASSBio-1911-treated astrocytes induces synapse formation suggests that LASSBio-1911 may induce the production of synaptogenic factors by astrocytes.

Altogether, our results suggest that the new iHDAC6 compound LASSBio-1911 induces synapse formation and protects against AβO-induced synapse loss through mitigation of inflammation and by inducing a shift in the astrocytic phenotype. Therefore, this study indicates that the modulation of acetylation through selective inhibition of HDAC6 in astrocytes may provide new insights and targets for the treatment of AD or other neurodegenerative diseases characterized by neuroinflammation, cognitive loss, and astrocyte reactivity.

## ACKNOWLEDGEMENTS

We thank Marcelo Meloni for technical assistance. We are indebted to Ricardo Lima-Filho for the preparation of Aβ oligomers.

## CONSENT STATEMENTS

No human consent was necessary since no human samples were used in the present work.

## REFERENCES

1. Hai Y, Christianson DW. Histone deacetylase 6 structure and molecular basis of catalysis and inhibition. Nature chemical biology. 2016;12:741–7.

2. Fischer A, Sananbenesi F, Wang X, Dobbin M, Tsai LH. Recovery of learning and memory is associated with chromatin remodelling. Nature. 2007;447:178–82.

3. Shukla S, Tekwani BL. Histone Deacetylases Inhibitors in Neurodegenerative Diseases, Neuroprotection and Neuronal Differentiation. Frontiers in pharmacology. 2020;11:537.

4. Rodrigues DA, Pinheiro PSM, Sagrillo FS, Bolognesi ML, Fraga CAM. Histone deacetylases as targets for the treatment of neurodegenerative disorders: Challenges and future opportunities. Medicinal research reviews. 2020;40:2177–211.

5. Ricobaraza A, Cuadrado-Tejedor M, Perez-Mediavilla A, Frechilla D, Del Rio J, Garcia-Osta A. Phenylbutyrate ameliorates cognitive deficit and reduces tau pathology in an Alzheimer’s disease mouse model. Neuropsychopharmacology: official publication of the American College of Neuropsychopharmacology. 2009;34:1721–32.

6. Kilgore M, Miller CA, Fass DM, Hennig KM, Haggarty SJ, Sweatt JD, et al. Inhibitors of class 1 histone deacetylases reverse contextual memory deficits in a mouse model of Alzheimer’s disease. Neuropsychopharmacology: official publication of the American College of Neuropsychopharmacology. 2010;35:870–80.

7. Kawaguchi Y, Kovacs JJ, McLaurin A, Vance JM, Ito A, Yao TP. The deacetylase HDAC6 regulates aggresome formation and cell viability in response to misfolded protein stress. Cell. 2003;115:727–38.

8. Kim C, Choi H, Jung ES, Lee W, Oh S, Jeon NL, et al. HDAC6 inhibitor blocks amyloid beta-induced impairment of mitochondrial transport in hippocampal neurons. PLoS One. 2012;7:e42983.

9. Choi H, Kim HJ, Yang J, Chae S, Lee W, Chung S, et al. Acetylation changes tau interactome to degrade tau in Alzheimer’s disease animal and organoid models. Aging Cell. 2020;19:e13081.

10. Ferreira ST, Klein WL. The Abeta oligomer hypothesis for synapse failure and memory loss in Alzheimer’s disease. Neurobiol Learn Mem. 2011;96:529–43.

11. Young-Pearse TL, Lee H, Hsieh YC, Chou V, Selkoe DJ. Moving beyond amyloid and tau to capture the biological heterogeneity of Alzheimer’s disease. Trends Neurosci. 2023;46:426–44.

12. Kumar A, Koistinen NA, Malarte ML, Nennesmo I, Ingelsson M, Ghetti B, et al. Astroglial tracer BU99008 detects multiple binding sites in Alzheimer’s disease brain. Molecular psychiatry. 2021;26:5833–47.

13. Orre M, Kamphuis W, Osborn LM, Jansen AHP, Kooijman L, Bossers K, et al. Isolation of glia from Alzheimer’s mice reveals inflammation and dysfunction. Neurobiol Aging. 2014;35:2746–60.

14. Diniz LP, Tortelli V, Matias I, Morgado J, Bergamo Araujo AP, Melo HM, et al. Astrocyte Transforming Growth Factor Beta 1 Protects Synapses against Abeta Oligomers in Alzheimer’s Disease Model. J Neurosci. 2017;37:6797–809.

15. Liddelow SA, Guttenplan KA, Clarke LE, Bennett FC, Bohlen CJ, Schirmer L, et al. Neurotoxic reactive astrocytes are induced by activated microglia. Nature. 2017;541:481–7.

16. Bellaver B, Povala G, Ferreira PCL, Ferrari-Souza JP, Leffa DT, Lussier FZ, et al. Astrocyte reactivity influences amyloid-beta effects on tau pathology in preclinical Alzheimer’s disease. Nature medicine. 2023.

17. Clarke LE, Liddelow SA, Chakraborty C, Munch AE, Heiman M, Barres BA. Normal aging induces A1-like astrocyte reactivity. Proc Natl Acad Sci U S A. 2018;115:E1896–E905.

18. Matias I, Diniz LP, Damico IV, Araujo APB, Neves LDS, Vargas G, et al. Loss of lamin-B1 and defective nuclear morphology are hallmarks of astrocyte senescence in vitro and in the aging human hippocampus. Aging Cell. 2022;21:e13521.

19. Matias I, Morgado J, Gomes FCA. Astrocyte Heterogeneity: Impact to Brain Aging and Disease. Front Aging Neurosci. 2019;11:59.

20. Rodrigues DA, Ferreira-Silva GA, Ferreira AC, Fernandes RA, Kwee JK, Sant’Anna CM, et al. Design, Synthesis, and Pharmacological Evaluation of Novel N-Acylhydrazone Derivatives as Potent Histone Deacetylase 6/8 Dual Inhibitors. J Med Chem. 2016;59:655–70.

21. Spanos F, Liddelow SA. An Overview of Astrocyte Responses in Genetically Induced Alzheimer’s Disease Mouse Models. Cells. 2020;9.

22. Mirzaei N, Davis N, Chau TW, Sastre M. Astrocyte Reactivity in Alzheimer’s Disease: Therapeutic Opportunities to Promote Repair. Curr Alzheimer Res. 2022;19:1–15.

23. Ye J, Zhong S, Deng Y, Yao X, Liu Q, Wang JZ, et al. HDAC7 Activates IKK/NF-kappaB Signaling to Regulate Astrocyte-Mediated Inflammation. Mol Neurobiol. 2022;59:6141–57.

24. Kanski R, Sneeboer MA, van Bodegraven EJ, Sluijs JA, Kropff W, Vermunt MW, et al. Histone acetylation in astrocytes suppresses GFAP and stimulates a reorganization of the intermediate filament network. J Cell Sci. 2014;127:4368–80.

25. Wu X, Chen PS, Dallas S, Wilson B, Block ML, Wang CC, et al. Histone deacetylase inhibitors up-regulate astrocyte GDNF and BDNF gene transcription and protect dopaminergic neurons. Int J Neuropsychopharmacol. 2008;11:1123–34.

26. Mooij WT, Verdonk ML. General and targeted statistical potentials for protein-ligand interactions. Proteins. 2005;61:272–87.

27. Korb O, Stutzle T, Exner TE. Empirical scoring functions for advanced protein-ligand docking with PLANTS. J Chem Inf Model. 2009;49:84–96.

28. Eldridge MD, Murray CW, Auton TR, Paolini GV, Mee RP. Empirical scoring functions: I. The development of a fast empirical scoring function to estimate the binding affinity of ligands in receptor complexes. J Comput Aided Mol Des. 1997;11:425–45.

29. Jones G, Willett P, Glen RC. Molecular recognition of receptor sites using a genetic algorithm with a description of desolvation. J Mol Biol. 1995;245:43–53.

30. De Felice FG, Velasco PT, Lambert MP, Viola K, Fernandez SJ, Ferreira ST, et al. Abeta oligomers induce neuronal oxidative stress through an N-methyl-D-aspartate receptor-dependent mechanism that is blocked by the Alzheimer drug memantine. J Biol Chem. 2007;282:11590–601.

31. De Felice FG, Wu D, Lambert MP, Fernandez SJ, Velasco PT, Lacor PN, et al. Alzheimer’s disease-type neuronal tau hyperphosphorylation induced by A beta oligomers. Neurobiol Aging. 2008;29:1334–47.

32. Sebollela A, Freitas-Correa L, Oliveira FF, Paula-Lima AC, Saraiva LM, Martins SM, et al. Amyloid-beta oligomers induce differential gene expression in adult human brain slices. The Journal of biological chemistry. 2012;287:7436–45.

33. Figueiredo CP, Clarke JR, Ledo JH, Ribeiro FC, Costa CV, Melo HM, et al. Memantine rescues transient cognitive impairment caused by high-molecular-weight abeta oligomers but not the persistent impairment induced by low-molecular-weight oligomers. J Neurosci. 2013;33:9626–34.

34. Diniz LP, Almeida JC, Tortelli V, Vargas Lopes C, Setti-Perdigao P, Stipursky J, et al. Astrocyte-induced synaptogenesis is mediated by transforming growth factor beta signaling through modulation of D-serine levels in cerebral cortex neurons. J Biol Chem. 2012;287:41432–45.

35. Guevara I, Iwanejko J, Dembinska-Kiec A, Pankiewicz J, Wanat A, Anna P, et al. Determination of nitrite/nitrate in human biological material by the simple Griess reaction. Clin Chim Acta. 1998;274:177–88.

36. Figueiredo CP, Ferreira NC, Passos GF, Costa R, Neves FS, Machado CS, et al. Toxicological Evaluation of Anti-Scrapie Trimethoxychalcones and Oxadiazoles. An Acad Bras Cienc. 2015;87:1421–34.

37. Ahn SY, Yoo HS, Lee JH, Sung DK, Jung YJ, Sung SI, et al. Quantitative in vivo detection of brain cell death after hypoxia ischemia using the lipid peak at 1.3 ppm of proton magnetic resonance spectroscopy in neonatal rats. J Korean Med Sci. 2013;28:1071–6.

38. Livak KJ, Schmittgen TD. Analysis of relative gene expression data using real-time quantitative PCR and the 2(-Delta Delta C(T)) Method. Methods. 2001;25:402–8.

39. Diniz LP, Matias I, Araujo APB, Garcia MN, Barros-Aragao FGQ, Alves-Leon SV, et al. alpha-synuclein oligomers enhance astrocyte-induced synapse formation through TGF-beta1 signaling in a Parkinson’s disease model. J Neurochem. 2019;150:138–57.

40. Finnin MS, Donigian JR, Cohen A, Richon VM, Rifkind RA, Marks PA, et al. Structures of a histone deacetylase homologue bound to the TSA and SAHA inhibitors. Nature. 1999;401:188–93.

41. Sofroniew MV. Astrocyte Reactivity: Subtypes, States, and Functions in CNS Innate Immunity. Trends Immunol. 2020;41:758–70.

42. Simoes-Pires C, Zwick V, Nurisso A, Schenker E, Carrupt PA, Cuendet M. HDAC6 as a target for neurodegenerative diseases: what makes it different from the other HDACs? Mol Neurodegener. 2013;8:7.

43. Govindarajan N, Rao P, Burkhardt S, Sananbenesi F, Schluter OM, Bradke F, et al. Reducing HDAC6 ameliorates cognitive deficits in a mouse model for Alzheimer’s disease. EMBO Mol Med. 2013;5:52–63.

44. Ceyzeriat K, Ben Haim L, Denizot A, Pommier D, Matos M, Guillemaud O, et al. Modulation of astrocyte reactivity improves functional deficits in mouse models of Alzheimer’s disease. Acta neuropathologica communications. 2018;6:104.

45. Subramanian J, Savage JC, Tremblay ME. Synaptic Loss in Alzheimer’s Disease: Mechanistic Insights Provided by Two-Photon in vivo Imaging of Transgenic Mouse Models. Front Cell Neurosci. 2020;14:592607.

46. Duclot F, Kabbaj M. The Role of Early Growth Response 1 (EGR1) in Brain Plasticity and Neuropsychiatric Disorders. Front Behav Neurosci. 2017;11:35.

47. Hampel H, Hardy J, Blennow K, Chen C, Perry G, Kim SH, et al. The Amyloid-beta Pathway in Alzheimer’s Disease. Mol Psychiatry. 2021;26:5481–503.

48. Benek O, Korabecny J, Soukup O. A Perspective on Multi-target Drugs for Alzheimer’s Disease. Trends Pharmacol Sci. 2020;41:434–45.

49. Guerra FS, Rodrigues DA, Fraga CAM, Fernandes PD. Novel Single Inhibitor of HDAC6/8 and Dual Inhibitor of PI3K/HDAC6 as Potential Alternative Treatments for Prostate Cancer. Pharmaceuticals (Basel). 2021;14.

50. Peleg S, Sananbenesi F, Zovoilis A, Burkhardt S, Bahari-Javan S, Agis-Balboa RC, et al. Altered histone acetylation is associated with age-dependent memory impairment in mice. Science. 2010;328:753–6.

51. Lu X, Wang L, Yu C, Yu D, Yu G. Histone Acetylation Modifiers in the Pathogenesis of Alzheimer’s Disease. Front Cell Neurosci. 2015;9:226.

52. Zhang K, Schrag M, Crofton A, Trivedi R, Vinters H, Kirsch W. Targeted proteomics for quantification of histone acetylation in Alzheimer’s disease. Proteomics. 2012;12:1261–8.

53. Francis YI, Fa M, Ashraf H, Zhang H, Staniszewski A, Latchman DS, et al. Dysregulation of histone acetylation in the APP/PS1 mouse model of Alzheimer’s disease. J Alzheimers Dis. 2009;18:131–9.

54. Hugais MM, Cobos SN, Bennett SA, Paredes J, Foran G, Torrente MP. Changes in Histone H3 Acetylation on Lysine 9 Accompany Abeta 1-40 Overexpression in an Alzheimer’s Disease Yeast Model. MicroPubl Biol. 2021;2021.

55. Janczura KJ, Volmar CH, Sartor GC, Rao SJ, Ricciardi NR, Lambert G, et al. Inhibition of HDAC3 reverses Alzheimer’s disease-related pathologies in vitro and in the 3xTg-AD mouse model. Proc Natl Acad Sci U S A. 2018;115:E11148–E57.

56. Krishna K, Behnisch T, Sajikumar S. Inhibition of Histone Deacetylase 3 Restores Amyloid-beta Oligomer-Induced Plasticity Deficit in Hippocampal CA1 Pyramidal Neurons. J Alzheimers Dis. 2016;51:783–91.

57. Onishi T, Maeda R, Terada M, Sato S, Fujii T, Ito M, et al. A novel orally active HDAC6 inhibitor T-518 shows a therapeutic potential for Alzheimer’s disease and tauopathy in mice. Sci Rep. 2021;11:15423.

58. Li Y, Sang S, Ren W, Pei Y, Bian Y, Chen Y, et al. Inhibition of Histone Deacetylase 6 (HDAC6) as a therapeutic strategy for Alzheimer’s disease: A review (2010-2020). Eur J Med Chem. 2021;226:113874.

59. Sabnis RW. Novel Histone Deacetylase 6 Inhibitors for Treating Alzheimer’s Disease and Cancer. ACS Med Chem Lett. 2021;12:1202–3.

60. Zhang Y, Li N, Caron C, Matthias G, Hess D, Khochbin S, et al. HDAC-6 interacts with and deacetylates tubulin and microtubules in vivo. EMBO J. 2003;22:1168–79.

61. Li Y, Shin D, Kwon SH. Histone deacetylase 6 plays a role as a distinct regulator of diverse cellular processes. FEBS J. 2013;280:775–93.

62. Carlomagno Y, Chung DC, Yue M, Castanedes-Casey M, Madden BJ, Dunmore J, et al. An acetylation-phosphorylation switch that regulates tau aggregation propensity and function. The Journal of biological chemistry. 2017;292:15277–86.

63. Boyault C, Zhang Y, Fritah S, Caron C, Gilquin B, Kwon SH, et al. HDAC6 controls major cell response pathways to cytotoxic accumulation of protein aggregates. Genes & development. 2007;21:2172–81.

64. Kalinin S, Polak PE, Lin SX, Braun D, Guizzetti M, Zhang X, et al. Dimethyl fumarate regulates histone deacetylase expression in astrocytes. J Neuroimmunol. 2013;263:13–9.

65. Patnala R, Arumugam TV, Gupta N, Dheen ST. HDAC Inhibitor Sodium Butyrate-Mediated Epigenetic Regulation Enhances Neuroprotective Function of Microglia During Ischemic Stroke. Mol Neurobiol. 2017;54:6391–411.

66. MacDonald JL, Roskams AJ. Histone deacetylases 1 and 2 are expressed at distinct stages of neuro-glial development. Dev Dyn. 2008;237:2256–67.

67. Faraco G, Pittelli M, Cavone L, Fossati S, Porcu M, Mascagni P, et al. Histone deacetylase (HDAC) inhibitors reduce the glial inflammatory response in vitro and in vivo. Neurobiol Dis. 2009;36:269–79.

68. Ding ZB, Song LJ, Wang Q, Kumar G, Yan YQ, Ma CG. Astrocytes: a double-edged sword in neurodegenerative diseases. Neural Regen Res. 2021;16:1702–10.

69. Katsouri L, Birch AM, Renziehausen AWJ, Zach C, Aman Y, Steeds H, et al. Ablation of reactive astrocytes exacerbates disease pathology in a model of Alzheimer’s disease. Glia. 2020;68:1017–30.

70. Jayakumar AR, Tong XY, Curtis KM, Ruiz-Cordero R, Shamaladevi N, Abuzamel M, et al. Decreased astrocytic thrombospondin-1 secretion after chronic ammonia treatment reduces the level of synaptic proteins: in vitro and in vivo studies. J Neurochem. 2014;131:333–47.

71. Hinkle JT, Dawson VL, Dawson TM. The A1 astrocyte paradigm: New avenues for pharmacological intervention in neurodegeneration. Mov Disord. 2019;34:959–69.

72. Yun SP, Kam TI, Panicker N, Kim S, Oh Y, Park JS, et al. Block of A1 astrocyte conversion by microglia is neuroprotective in models of Parkinson’s disease. Nat Med. 2018;24:931–8.

73. Park JS, Kam TI, Lee S, Park H, Oh Y, Kwon SH, et al. Blocking microglial activation of reactive astrocytes is neuroprotective in models of Alzheimer’s disease. Acta Neuropathol Commun. 2021;9:78.

74. Liu L, Zhou X, Shetty S, Hou G, Wang Q, Fu J. HDAC6 inhibition blocks inflammatory signaling and caspase-1 activation in LPS-induced acute lung injury. Toxicol Appl Pharmacol. 2019;370:178–83.

75. Park JK, Shon S, Yoo HJ, Suh DH, Bae D, Shin J, et al. Inhibition of histone deacetylase 6 suppresses inflammatory responses and invasiveness of fibroblast-like-synoviocytes in inflammatory arthritis. Arthritis Res Ther. 2021;23:177.

76. Diniz LP, Matias I, Siqueira M, Stipursky J, Gomes FCA. Astrocytes and the TGF-beta1 Pathway in the Healthy and Diseased Brain: a Double-Edged Sword. Mol Neurobiol. 2019;56:4653–79.

77. Buosi AS, Matias I, Araujo APB, Batista C, Gomes FCA. Heterogeneity in Synaptogenic Profile of Astrocytes from Different Brain Regions. Mol Neurobiol. 2018;55:751–62.

78. Allen NJ, Eroglu C. Cell Biology of Astrocyte-Synapse Interactions. Neuron. 2017;96:697–708.

79. Laskowski RA, Swindells MB. LigPlot+: multiple ligand-protein interaction diagrams for drug discovery. Journal of chemical information and modeling. 2011;51:2778–86.

80. Wallace AC, Laskowski RA, Thornton JM. LIGPLOT: a program to generate schematic diagrams of protein-ligand interactions. Protein engineering. 1995;8:127–34.

